# A novel truncated CHAP modular endolysin, CHAP^SAP26^-161, that lyses *Staphylococcus aureus*, *Acinetobacter baumannii,* and *Clostridioides difficile*

**DOI:** 10.1101/2024.02.22.581501

**Authors:** Yoon-Jung Choi, Shukho Kim, Ram Hari Dahal, Jungmin Kim

## Abstract

**Background:** Development of novel antimicrobial agents is imperative due to the increasing threat of antibiotic-resistant pathogens. This study aimed to validate the enhanced antibacterial activity and *in vivo* efficacy of a novel truncated endolysin, CHAP^SAP26^-161, derived from the CHAP domain of LysSAP26, against multidrug-resistant bacteria.

**Methods:** Two deletion mutants, CHAP^SAP26^-139 and CHAP^SAP26^-161, were constructed by deleting the C-terminal portion of LysSAP26. These were cloned and expressed, and their antibacterial activities, together with protein purification efficiency, were evaluated against 12 bacterial species under various environmental conditions. To test the temperature and pH stability of the three recombinant proteins, the antibacterial effects of the proteins at various temperatures (4°C–60°C) and pH values (3–10) were measured. Time-kill assay measured the optical density (600 nm) and colony-forming units after incubation for 0, 2, 4, 6, 8, and 24 h. We verified this through *in vivo* experiments using mouse models to evaluate the therapeutic potential of CHAP^SAP26^-161 against *Acinetobacter baumannii*.

**Results:** CHAP^SAP26^-161 exhibited higher protein purification efficiency and antibacterial activity than LysSAP26. Moreover, CHAP^SAP26^-161 showed the highest lytic activity against *A. baumannii* with a minimal bactericidal concentration (MBC) of 5–10 µg/mL, followed by *Staphylococcus aureus* with an MBC of 10–25 µg/mL. Interestingly, CHAP^SAP26^-161 could lyse anaerobic bacteria, such as *C. difficile*, with an MBC of 25–50 µg/mL. At pH 4–8 and temperatures of 4°C–45°C, CHAP^SAP26^-161 exhibited optimal hydrolase activity. The lytic activity of CHAP^SAP26^-161 was dependent on divalent metal ions, especially Zn^2+^, and increased in the presence of ethylenediamine tetraacetic acid. CHAP^SAP26^-161 demonstrated superior protein purification efficiency and antibacterial activity than LysSAP26. It showed high lytic activity against gram-positive, gram-negative, and anaerobic bacteria, including *S. aureus* and *Clostridioides difficile.* Enhanced stability under varied temperatures and pH conditions. *In vivo,* tests demonstrated promising therapeutic effects of CHAP^SAP26^-161 in murine systemic *A. baumannii* infection models.

**Conclusions:** CHAP^SAP26^-161, a truncated modular endolysin containing only the CHAP domain of LysSAP26, demonstrated higher protein purification efficiency and antibacterial activity than LysSAP26. It also exhibited extended-spectrum antibacterial activity against gram-positive, gram-negative, and anaerobic bacteria, such as *S. aureus*, *A. baumannii*, and *C. difficile*. Its successful *in vivo* application in murine models highlights its potential as an alternative therapeutic agent in combating antibiotic resistance.

## Introduction

Increasing prevalence of antibiotic-resistant bacteria, including multidrug-resistant (MDR), extensively drug-resistant (XDR), and pandrug-resistant strains, presents a formidable challenge in contemporary medical practice (Abram et al., 2020; Boneca, 2021; MC et al., 2021; Ozma et al., 2022; Souza et al., 2021). Therefore, there is an urgent need to develop alternative therapeutic strategies to combat these pathogens. Bacteriophages (phages) and their derivative endolysins have emerged as promising candidates in this endeavor (Fujimoto and Uematsu, 2022; Khan and Joshi, 2022; Kortright et al., 2019; Mondal et al., 2021, 2020; Sekiya et al., 2022; Ul Haq et al., 2012). Endolysins, hydrolytic enzymes produced by phages, demonstrate a broad host spectrum and possess the advantages of rapid bacterial cell lysis, low risk of resistance development, and efficacy against biofilms and mucosal surfaces (Abdelrahman et al., 2021; Borysowski et al., 2006; Gondil et al., 2020; Heselpoth et al., 2021; Mirski et al., 2019; Nelson et al., 2012; Schmelcher et al., 2012; Schmelcher and Loessner, 2016). The structure of bacteriophage endolysins varies, typically comprising an N-terminal catalytic domain and a C-terminal cell wall binding domain (DC et al., 2012; Stone et al., 2019). The catalytic domain cleaves the peptidoglycan layer of the bacterial cell wall, whereas the cell wall binding domain ensures specificity to target bacterial species (Becker et al., 2015; Filatova et al., 2010; Kretzer et al., 2007). The cysteine (Cys)–histidine (His)-dependent amidohydrolases/peptidases (CHAP) domain, a component of the catalytic module of endolysins, plays a critical role in antibacterial activity by hydrolyzing peptidoglycan bonds in the bacterial cell wall (Bateman and Rawlings, 2003; Becker et al., 2015; Fenton et al., 2011; Stacy et al., 2021; Sundarrajan et al., 2014; Vollmer et al., 2008). This domain, characterized by a conserved Cys–His motif and a distinctive structural fold, is often coupled with other functional domains to enhance antibacterial efficacy.

Our previous research reported on LysSAP26 (Kim et al., 2020b), consisting of 251 amino acids and a CHAP domain from amino acids 20–109 and an unknown functional protein from amino acids 110–251. The present study aimed to modify LysSAP26 to improve its antibacterial activity and protein purification efficiency. By deleting the C-terminal portion of LysSAP26, we produced two deletion mutants, CHAP^SAP26^-139 and CHAP^SAP26^-161, and compared their efficacy with the original protein. This decision was guided by prior studies showing the significant role of the CHAP domain in endolysins, such as LysK, and the impact of C-terminal truncations on bactericidal activity, as observed in LysSAP33 (Filatova et al., 2010; Horgan et al., 2009a; Kim et al., 2020b; O’Flaherty et al., 2005; Yu et al., 2021).

This study introduces CHAP^SAP26^-161, a novel truncated endolysin derived from the CHAP domain of LysSAP26 and explores its enhanced antibacterial activity. Our research underscores the broad-spectrum antibacterial potential of CHAP^SAP26^-161 and its promising application in treating multidrug-resistant *Acinetobacter baumannii* and *Staphylococcus aureus* infections, which are a growing concern in healthcare settings worldwide.

## Materials and Methods

### Bacterial strains and culture conditions

A comprehensive collection of 96 MDR clinical isolates, including *S. aureus*, *A. baumannii, Clostridioides difficile*, *Enterococcus faecium*, *Klebsiella pneumoniae*, *Escherichia coli*, *Pseudomonas aeruginosa*, *Enterococcus faecalis*, and other strains, was acquired from the Kyungpook National University Hospital Culture Collection for Pathogens. In addition, reference strains were acquired from the Korean Collection for Type Cultures and American Type Culture Collection (ATCC). The bacteria were cultured in appropriate media, such as Mueller–Hinton broth (DB, USA), Mueller–Hinton agar (MHA, DB, USA), brain heart infusion (BHI) broth, and blood agar plates, and incubated for 24–48 h at 37°C. Under specific conditions (5% H_2_, 5% CO_2_, and 90% N_2_), anaerobic bacteria, such as *C. difficile*, were cultured in an anaerobic chamber (Noor and Khetarpal, 2023; Sandhu and McBride, 2018). For long-term storage of the bacterial strains, BHI broth containing 15% glycerol (v/v) was used.

### Construction of chimeric endolysin expression vectors

The LysSAP26 sequence was used as a template to generate protein expression vectors for CHAP^SAP26^-139 and CHAP^SAP26^-161. PCR amplification involved specific primers (Supplementary Table 1) and Ex Taq Polymerase (Takara, Japan) (Choi et al., 2022; Kim et al., 2021, 2020b, 2020c, 2020a). The PCR products were then purified using a clean-up kit (GeneAll, Korea) and digested with NdeI and XhoI at 37°C for 1 h. The digested products were ligated into the pET-21a (+) vector using T4 DNA ligase at 18°C for 3 h. The ligation product was transformed into *E. coli* DH5α, and the transformed bacteria were selected on Luria–Bertani (LB) medium (BioShop, Canada) agar plates containing ampicillin (150 µg/mL). To amplify the insert to confirm successful cloning using the nde1-SAPlys primer and the T7 terminator primer, colony PCR was conduced (Choi et al., 2022; Kim et al., 2021, 2020b, 2020a, 2020c). Finally, the PCR products were sequenced to confirm the accuracy of the constructed vector.

### Expression and purification of endolysins

Following protocols detailed in references, each of the LysSAP26 (753 bp), CHAP^SAP26^-139 (417 bp), and CHAP^SAP26^-161 (483 bp) plasmids was transformed into *E. coli* BL21 (DE3) Star® cells (Choi et al., 2022; Kim et al., 2020b). Subsequent cultivation involved incubating the transformed cells in 1 L of LB medium supplemented with 150 μg/mL ampicillin at 37°C with a shaking speed of 150 rpm. This incubation continued until the optical density (OD) at 600 nm reached 0.5. To achieve a final concentration of 0.1 mM, isopropyl β-D-1-thiogalactopyranoside was then added followed by extended incubation at 18°C for 16 h. After incubation, cells were collected via centrifugation, resuspended in lysis buffer (50 mM Tris-HCl, 500 mM NaCl, and 1 mM ZnCl_2_; pH 7.4), and subsequently lysed using ultrasonication. The resulting supernatant was further processed via centrifugation and filtration and finally purified using a 5-mL His-trap affinity column. To eliminate residual imidazole from the purified fractions, dialysis was performed against the lysis buffer. The purity and molecular weights of the endolysins were verified via sodium dodecyl sulfate-polyacrylamide gel electrophoresis (SDS-PAGE) and western blot analysis. An anti-His-Tag monoclonal antibody (Ab Frontier, Korea) was used as the primary antibody to detect His-tagged proteins through western blotting, whereas a horseradish peroxidase-conjugated polyclonal rabbit antimouse immunoglobulin G (Dako, Denmark) was used as the secondary antibody. The concentration of the purified proteins was quantified using the Pierce BCA Protein Assay Kit (Thermo Fisher Scientific, Waltham, MA, United States) as described previously (Choi et al., 2022; Kim et al., 2020b).

### Bioinformatic analysis

To elucidate the physical, chemical, and structural characteristics of the CHAP^SAP26^-139 and CHAP^SAP26^-161 protein sequences, a thorough bioinformatic examination was conducted (refer to Figure 1A). The alignment and validation of these sequences were executed using the BLASTP and Clustal Omega algorithms. Expasy’s ProtParam tool (https://web.expasy.org/protparam/) was used to predict various key properties, such as molecular weight, theoretical isoelectric point (pI), and extinction coefficient. In addition, the i-TASSER unified platform (https://seq2fun.dcmb.med.umich.edu/I-TASSER/) was used to analyze the three-dimensional structures and potential functional attributes of these proteins.

**Figure 1.**
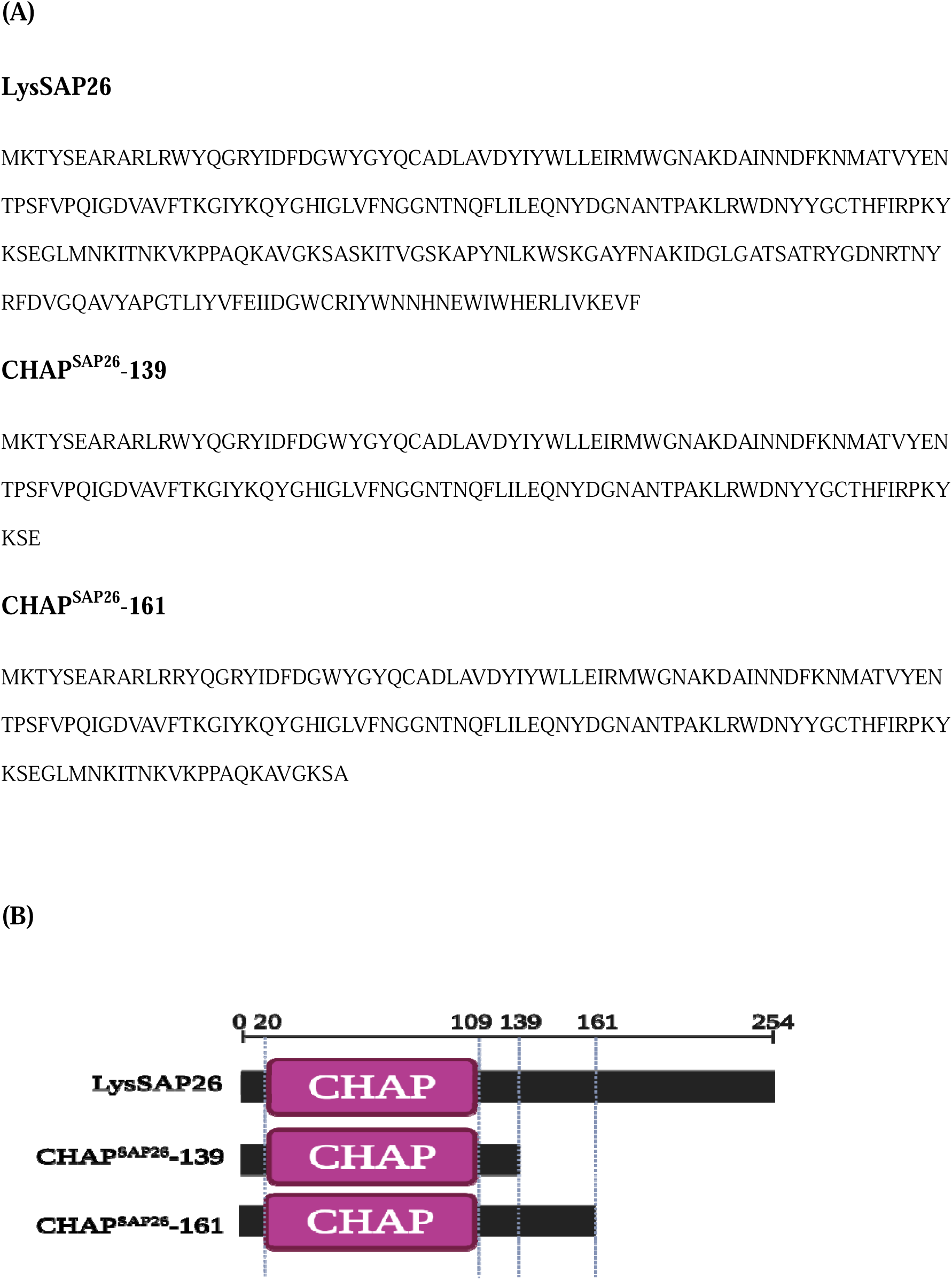
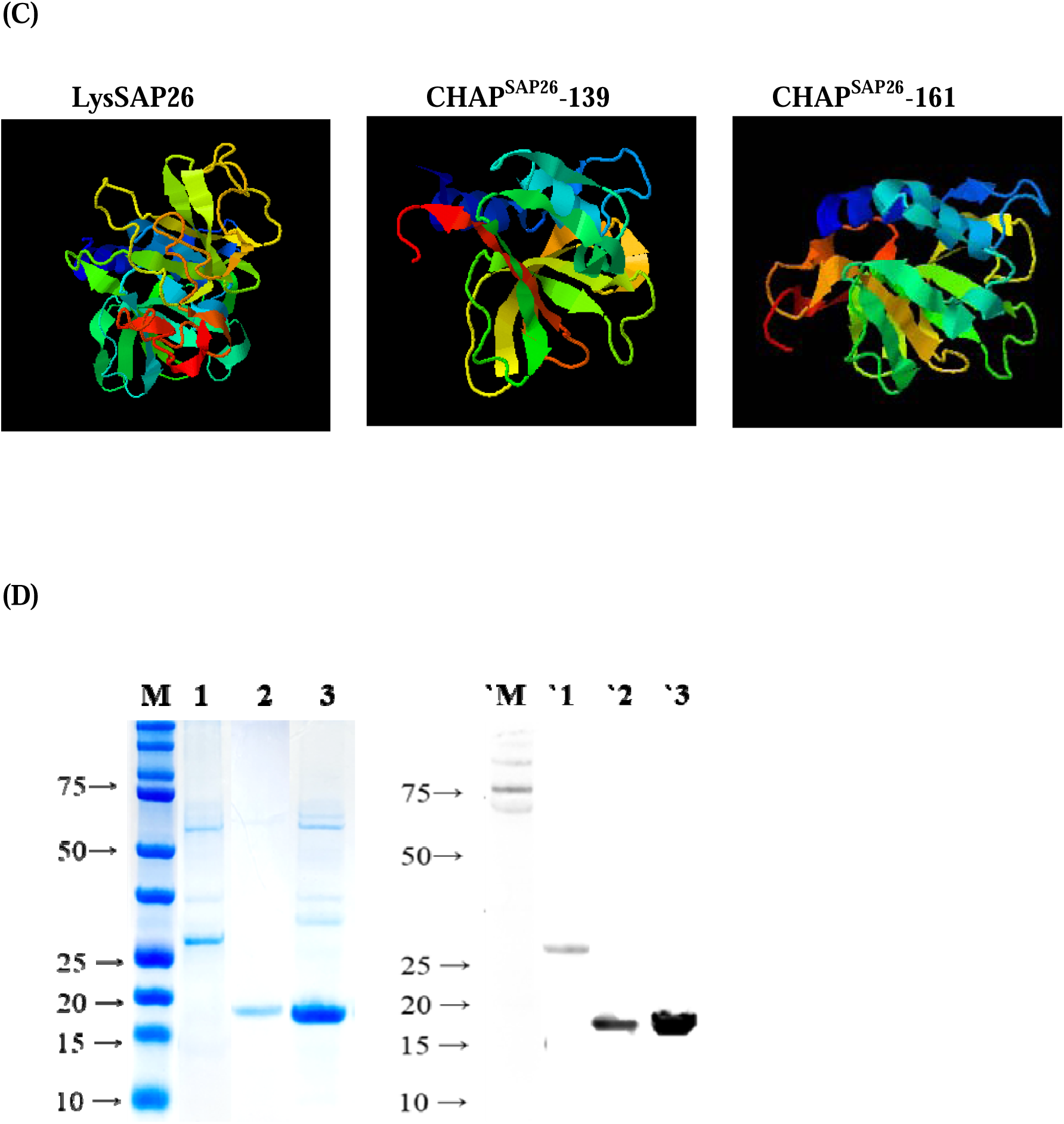
Structural characterization of endolysins. (A) Comparative amino acid sequence analysis of LysSAP26, CHAP^SAP26^-139, and CHAP^SAP26^-161; (B) Schematic representation of the domain architecture for the three endolysins; (C) Visualization of the three-dimensional structures of the endolysins, analyzed using the i-TASSER unified platform to evaluate potential functional characteristics of these proteins; (D) Protein size comparison through SDS-PAGE and western blotting, featuring LysSAP26 (29.1 kDa), CHAP^SAP26^-161 (18.6 kDa), and CHAP^SAP26^-139 (16.2 kDa).

### Determination of the antibacterial activity of endolysins

Using a modified broth microdilution method in 96-well, round-bottomed microplates (SPL, Korea), the minimal inhibitory concentration (MIC) and MBC of the endolysins were determined according to the CLSI guidelines (CLSI, 2020; “CLSI GUIDELINES 2020,” 2020). Various concentrations (1–100 μg/mL) of endolysins were added to the bacterial cells (10^6^ CFU/mL) and incubated for 18–24 h at 37°C. MIC was identified as the lowest endolysin concentration that completely inhibited bacterial growth. A 10-μL sample from each well of the MIC assay was transferred onto MHA plates to determine the MBC, followed by further incubation for 18–24 h to establish bactericidal activity. All assays were performed in triplicate to ensure reproducibility and reliability of the results.

### Time-kill assay

A time-kill analysis was conducted to elucidate the time-sustained antimicrobial efficacy of CHAP^SAP26^-161 against *A. baumannii* ATCC 17978 and *S. aureus* ATCC 25923(Kim et al., 2020b). For this purpose, bacterial suspensions were prepared using the microdilution method, and CHAP^SAP26^-161 was added to achieve final concentrations of 10–50 µg/mL. The suspensions were incubated for 0, 2, 4, 6, 8, and 24 h, after which they were plated on LB agar to determine the CFUs. A dialysis buffer was used as a control instead of CHAP^SAP26^-161. To ensure reproducibility of the results, the experiment was conducted twice.

### Impact of physicochemical conditions on the enzymatic activity of endolysins

To evaluate the influence of various physicochemical conditions on the antimicrobial activity of endolysins, turbidity reduction assays were used. These assays specifically explored the effects of a broad temperature range (4°C to 60°C), pH levels ranging from 3 to 10, and different concentrations of NaCl (up to 500 mM). Furthermore, the assays assessed the impact of divalent metal ions, such as 1 mM CaCl_2_, CuCl_2_, MgCl_2_, ZnCl_2_, and ZnSO_4_, including the influence of EDTA concentrations up to 10 mM. The antibacterial activity of endolysins was determined after a 2-h incubation at each designated temperature. For pH stability assessment, the buffer pH was adjusted using NaOH and HCl before endolysin dialysis to evaluate enzyme stability under varying pH conditions. During dialysis, NaCl, divalent ions, and EDTA were incorporated at specified concentrations, after which the endolysin buffer was replaced for further analysis. To evaluate the impact of temperature, pH, NaCl concentration, divalent metal ions, and EDTA on the antimicrobial activity of endolysins, these turbidity reduction assays were also conducted. *A. baumannii* ATCC 17978 and *S. aureus* ATCC 25923 strains were incubated with endolysins at concentrations of 10–50 µg/mL under varying conditions for 18 h. The optical density at OD_600 nm_ of each well was measured to quantify antimicrobial activity. Statistical analyses were conducted to identify significant differences when compared with either the most active sample (e.g., optimal temperature, pH, and NaCl concentration) or the blank control (e.g., metal ion exposure).

### Assessment of the cytotoxic impact of CHAP^SAP26^-161 on human lung epithelial cells

Human lung epithelial cells (A549), which originate from adenocarcinoma, were used to evaluate the cytotoxic potential of CHAP^SAP26^-161. Cytotoxicity was assessed using the MTT assay (3-[4,5-methylthiazol-2-yl]-2,5-diphenyl-tetrazolium bromide, Amresco, Solon, OH, USA), following the manufacturer’s protocol. Initially, A549 cells (1 × 10^5^ cells/well) were plated in a 24-well plate with RPMI medium and incubated overnight in a CO_2_ incubator to allow for cell attachment. Subsequently, the medium was replaced with fresh RPMI containing varying concentrations of CHAP^SAP26^-161 (ranging from 25 to 1,000 μg/mL). Under these conditions, the cells were incubated for an additional 24 h. Following this period, the medium was discarded and the cells were rinsed with phosphate-buffered saline (PBS). Each well then received 250 μL of MTT solution (0.5 mg/mL), and the cells were further incubated for 2 h. Finally, 250 μL of a solubilizing solution (90% isopropanol, 0.01% Triton X-100, and 0.01N HCl) was added. The resultant color development, which is indicative of cell viability, was quantified at 570 nm using a VersaMax™ microplate reader (Molecular Devices, San Jose, CA, USA).

### Evaluation of the efficacy of CHAP^SAP26^-161 in *Acinetobacter baumannii* mouse models

*A. baumannii* ATCC 17978 was selected for the protection assay in a murine systemic infection model. Pathogen-free female BALB/c mice (6 weeks old, weight 16–19 g) were obtained from OrientBio (OrientBio, Seongnam, Korea). Neutropenia in mice was induced via intraperitoneal injection of cyclophosphamide (150 mg/kg) on days 4 and 1 before bacterial injection. Systemic infection was induced via intraperitoneal injection of 200 μL of a log-phase bacterial inoculum (1 × 10^9^ CFU). Mice were divided into six groups as follows (five mice/group):

- Group 1: Inactive control (200 μL PBS + 200 μL buffer A)
- Group 2: Infection control (200 μL *A. baumannii* + 200 μL buffer A)
- Group 3: CHAP^SAP26^-161 safety test (200 μL PBS + 50 μg/200 μL CHAP^SAP26^-161)
- Group 4: Positive control with LysSAP26 treatment (200 μL *A. baumannii* + 50 μg/200 μL LysSAP26)
- Group 5: CHAP^SAP26^-161 treatment (200 μL *A. baumannii* + 50 μg/200 μL CHAP^SAP26^-161)

Mice in each group were treated 30 min after infection, and postinfection survival was monitored once a day for 7 days following infection.

### Statistical analysis

Statistical analysis was conducted using OriginPro, applying one-way analysis of variance (ANOVA) followed by Tukey’s test for all pairwise comparisons (95% confidence interval) (Hecke, 2013; McHugh, 2011). Data are expressed as mean values with standard deviations, and statistical significance was set at a p value of <0.01 (Murdoch et al., 2012).

### Results Bioinformatics analysis and purification of endolysins

Two deletion mutants, CHAP^SAP26^-161 and CHAP^SAP26^-139, were engineered by truncating the C-terminal portion of the LysSAP26-coding gene (Figure 1A-1C). These constructs were successfully incorporated into the pET-21a (+) vector and expressed in *E. coli* cells. The molecular weights of the three purified proteins, LysSAP26 at 29.1 kDa, CHAP^SAP26^-161 at 18.6 kDa, and CHAP^SAP26^-139 at 16.2 kDa, were confirmed to match the expected values, as verified through SDS-PAGE (Figure 1D). The predicted PI values for CHAP^SAP26^-161 and CHAP^SAP26^-139 were 9.32 and 7.70, respectively. The protein concentrations were 2.67 mg/L for LysSAP26, 17.32 mg/L for CHAP^SAP26^-161, and 6.33 mg/L for CHAP^SAP26^-139 (Figure 1D).

### Antimicrobial activity of endolysins

The antibacterial activity of CHAP^SAP26^-161 and CHAP^SAP26^-139 was compared with that of LysSAP26 (Table 1). CHAP^SAP26^-161 showed a higher inhibitory activity than SAP26 against all bacterial species tested, whereas CHAP^SAP26^-139 did not. CHAP^SAP26^-161 exhibited two-to five-fold lower MIC and MBC than LysSAP26. CHAP^SAP26^-161 showed the highest lytic activity against *A. baumannii* with an MBC of 5–10 µg/mL, followed by *S. aureus* with an MBC of 10–25 µg/mL, among the 12 types of bacterial species tested. Interestingly, CHAP^SAP26^-161 demonstrated lytic activity against anaerobic bacteria, such as *C. difficile*, with an MBC of 25–50 µg/mL (Table 1), but not against other anaerobic bacteria, such as *C. acnes* and *F. varium* (Table 1).

**Table 1.**
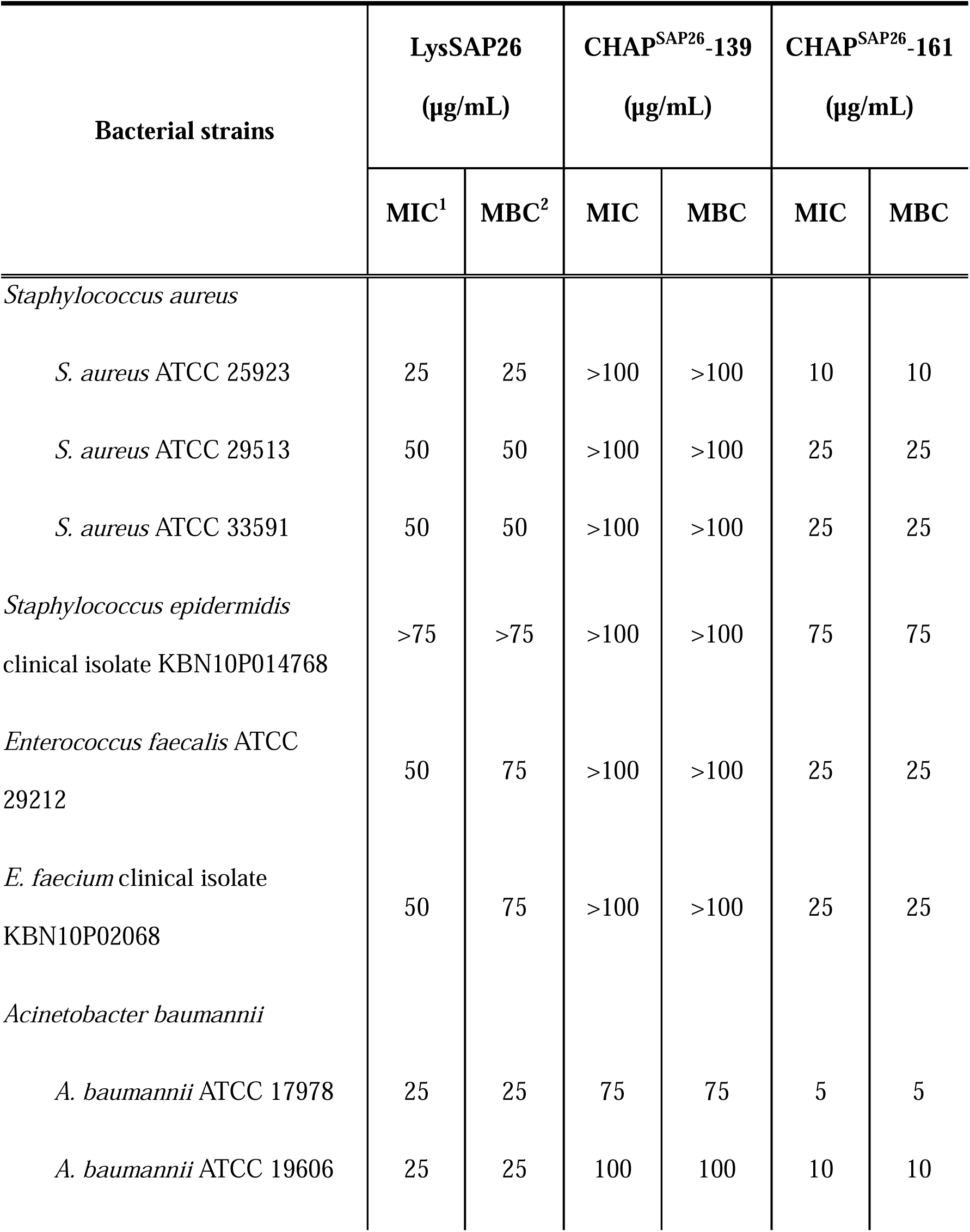
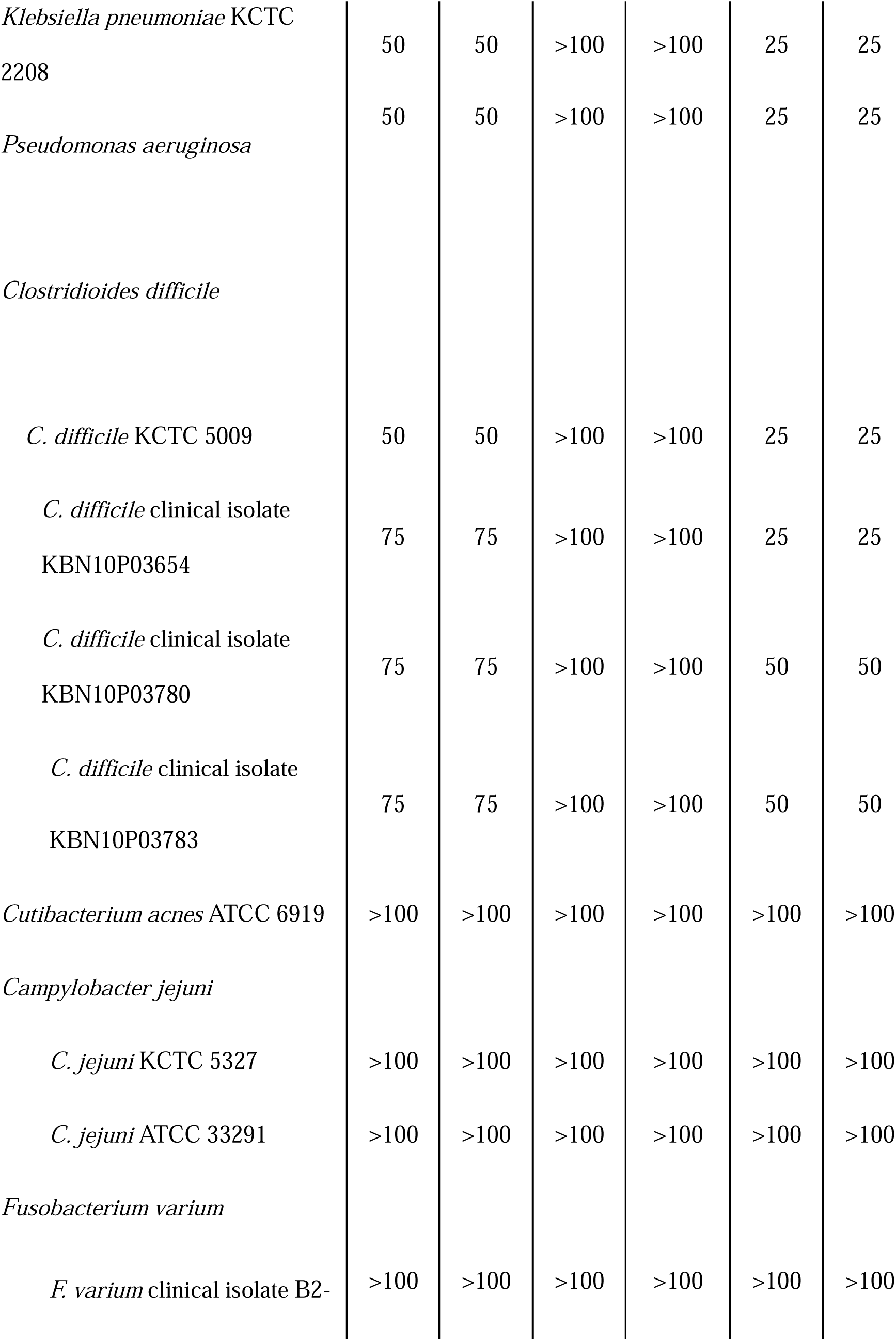
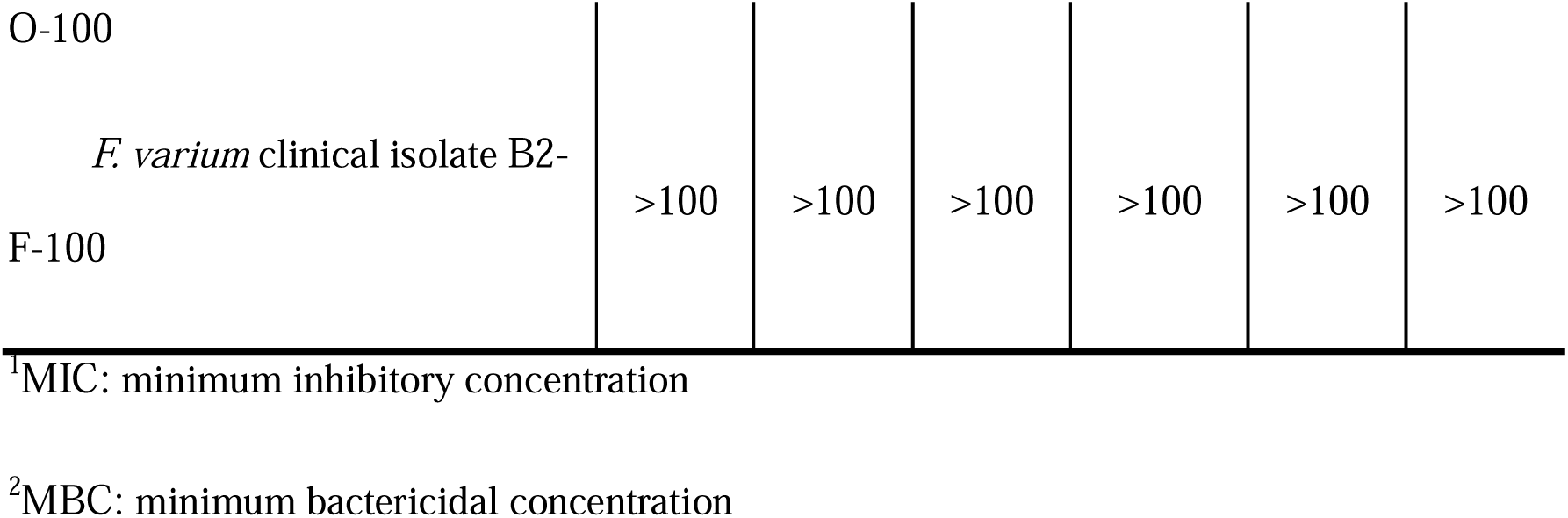
Antibacterial activity of LysSAP26, CHAP^SAP26^-161, and CHAP^SAP26^-139 against 18 bacterial strains.

Treatment with 25 µg/mL LysSAP26 or 5 µg/mL CHAP^SAP26^-161 decreased the CFU counts of *A. baumannii* that began to occur after 6 h and was maintained until 24 h (Figure 2A) during the time-kill assay. When *S. aureus* was treated with 25 µg/mL LysSAP26 or 10 µg/mL CHAP^SAP26^-161, the CFU counts, which began to decrease immediately after treatment, were 2 log CFU lower than the initial count after 12 h (Figure 2B).

**Figure 2.**
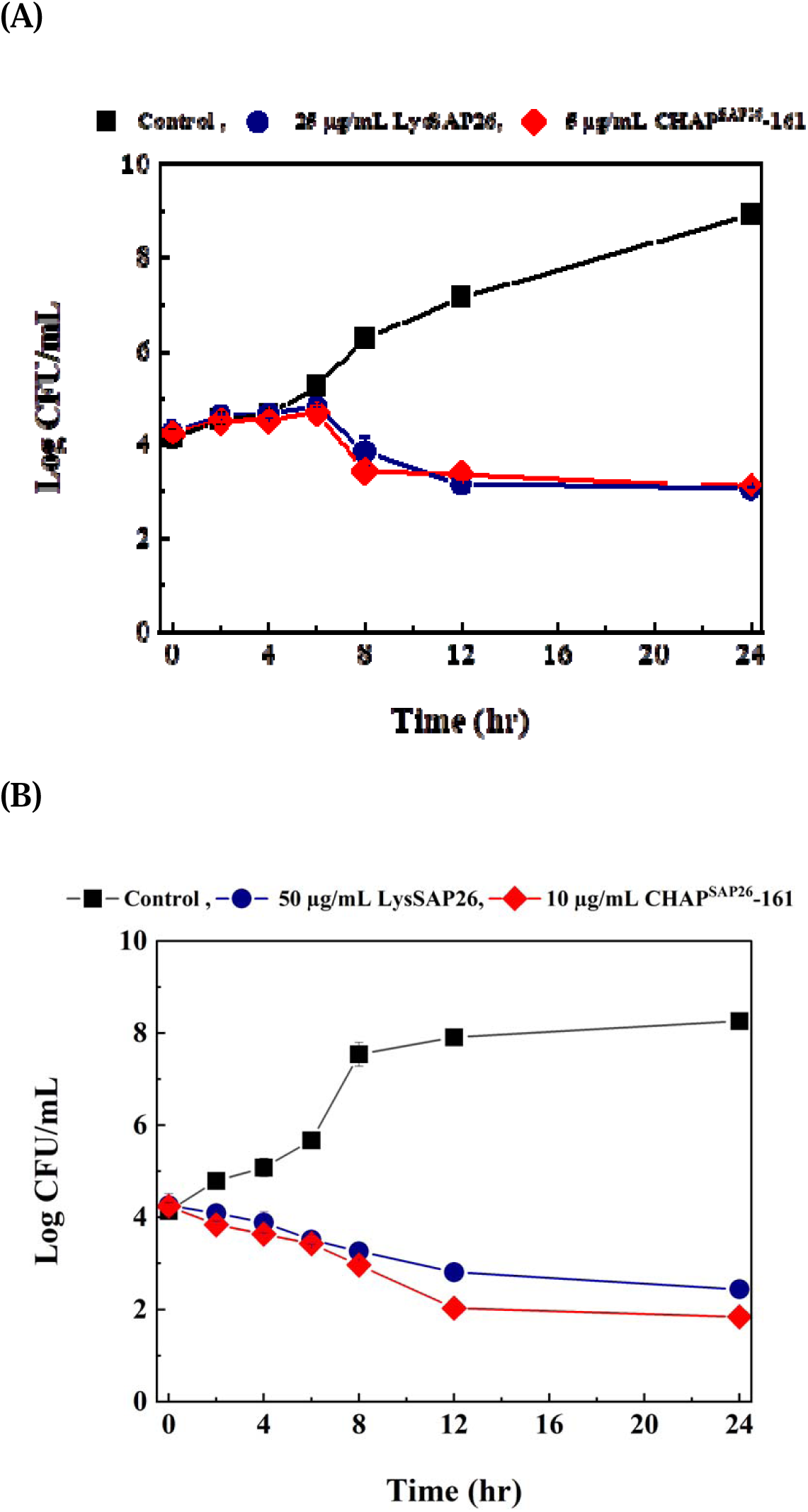
Time-dependent bactericidal effect of endolysins on *Acinetobacter baumannii* and *Staphylococcus aureus.* (A) Bactericidal effect against *A. baumannii* ATCC 17978 by LysSAP26 (25 μg/mL) and CHAP^SAP26^-161 (5 μg/mL); (B) Bactericidal effect against *S. aureus* ATCC 25923 by LysSAP26 (50 μg/mL) and CHAPSAP26-161 (10 μg/mL).

The antibacterial efficacy of LysSAP26 and CHAP^SAP26^-161 against clinical MDR strains was also compared. In total, 117 clinical isolates of *E. faecium*, *S. aureus*, *K. pneumoniae*, *A. baumannii*, *P. aeruginosa*, and *E. coli* were included in the susceptibility test, and the results are demonstrated in Table 2. Similar to the results presented in Table 1, CHAP^SAP26^-161 showed two-to five-fold lower MIC and MBC than LysSAP26 (Table 2). CHAP^SAP26^-161 demonstrated an MIC and MBC of 5–25 µg/mL against all 17 carbapenem-resistant *A. baumannii* isolates. The MIC values of CHAP^SAP26^-161 were 20 µg/mL for all 20 oxacillin-resistant *S. aureus* isolates, 25–50 µg/mL for 20 carbapenem-resistant *K. pneumoniae* isolates, and 25–50 µg/mL for 20 carbapenem-resistant *P. aeruginosa* isolates. Among 20 carbapenem-resistant or cephalosporin-resistant *E. coli* isolates, the MIC values were 50 µg/mL for 2 isolates and 25 µg/mL for 18 isolates. The MIC values of CHAP^SAP26^-161 for 20 vancomycin-resistant *E. faecium* isolates were 25 µg/mL (14 isolates) and 50 µg/mL (6 isolates) (Table 2).

**Table 2.**
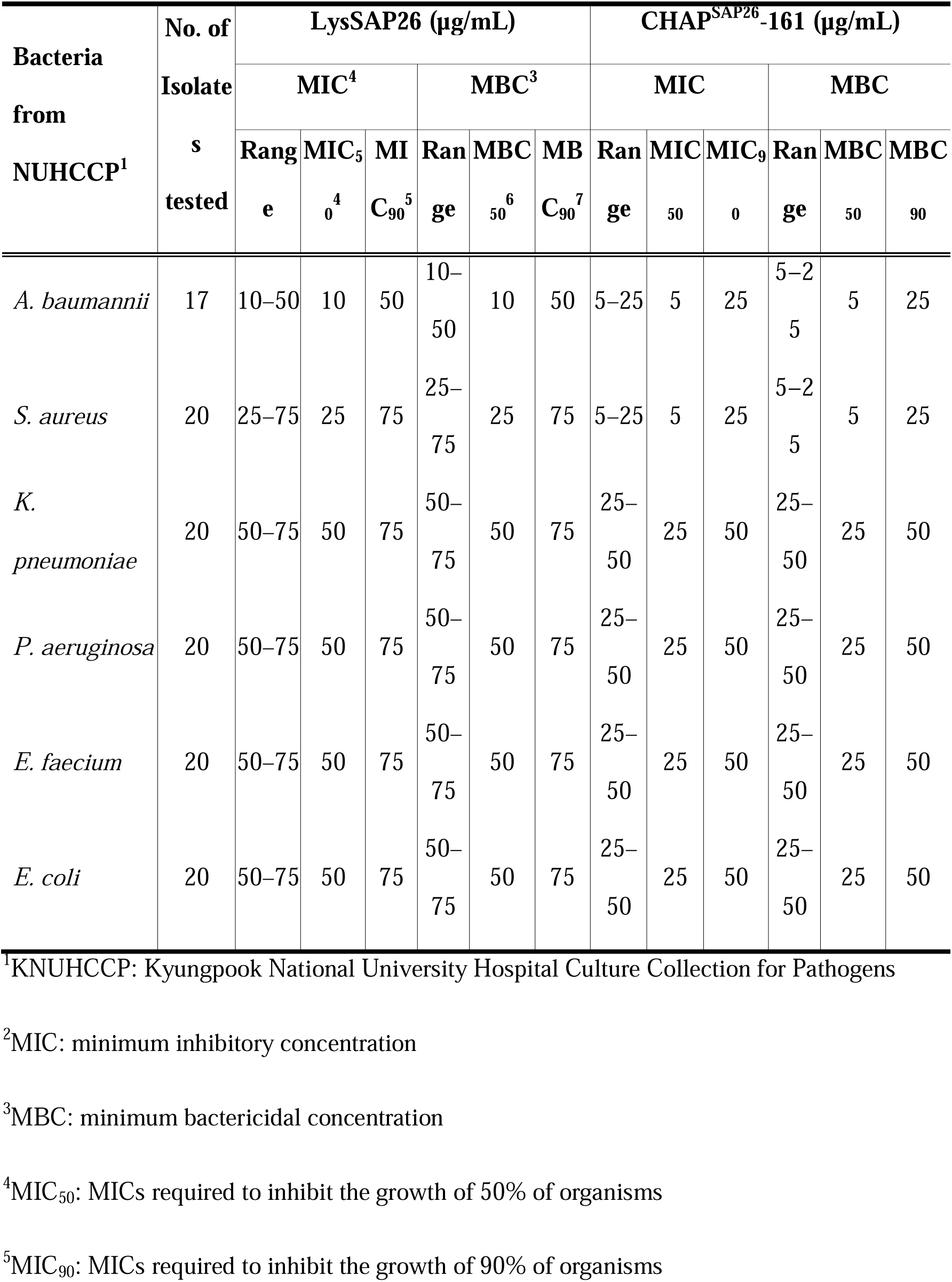
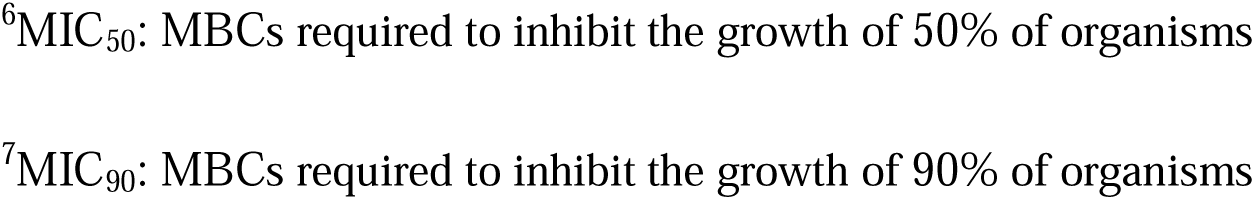
Antibacterial activity of LysSAP26 and CHAP^SAP26^-161 against MDR clinical isolates of ESKAPE pathogens.

| Bacteria<br>from<br>NUHCCP<sup>1</sup> | No. of<br>Isolate<br>s<br>tested | LysSAP26 (µg/mL) |  |  |  |  |  | CHAP<sup>SAP26</sup>-161 (µg/mL) |  |  |  |  |  |
| --- | --- | --- | --- | --- | --- | --- | --- | --- | --- | --- | --- | --- | --- |
|  |  | MIC<sup>4</sup> |  |  | MBC<sup>3</sup> |  |  | MIC |  |  | MBC |  |  |
|  |  | Rang<br>e | MIC<sub>5</sub><br>0<sup>4</sup> | MI<br>C<sub>90</sub><sup>5</sup> | Ran<br>ge | MBC<br>50<sup>6</sup> | MB<br>C<sub>90</sub><sup>7</sup> | Ran<br>ge | MIC<br>50 | MIC<sub>90</sub><br>0 | Ran<br>ge | MBC<br>50 | MBC<br>90 |
| *A. baumannii* | 17 | 10–50 | 10 | 50 | 10–<br>50 | 10 | 50 | 5–25 | 5 | 25 | 5–2<br>5 | 5 | 25 |
| *S. aureus* | 20 | 25–75 | 25 | 75 | 25–<br>75 | 25 | 75 | 5–25 | 5 | 25 | 5–2<br>5 | 5 | 25 |
| *K.*<br>*pneumoniae* | 20 | 50–75 | 50 | 75 | 50–<br>75 | 50 | 75 | 25–<br>50 | 25 | 50 | 25–<br>50 | 25 | 50 |
| *P. aeruginosa* | 20 | 50–75 | 50 | 75 | 50–<br>75 | 50 | 75 | 25–<br>50 | 25 | 50 | 25–<br>50 | 25 | 50 |
| *E. faecium* | 20 | 50–75 | 50 | 75 | 50–<br>75 | 50 | 75 | 25–<br>50 | 25 | 50 | 25–<br>50 | 25 | 50 |
| *E. coli* | 20 | 50–75 | 50 | 75 | 50–<br>75 | 50 | 75 | 25–<br>50 | 25 | 50 | 25–<br>50 | 25 | 50 |

<sup>1</sup>KNUHCCP: Kyungpook National University Hospital Culture Collection for Pathogens
<sup>2</sup>MIC: minimum inhibitory concentration
<sup>3</sup>MBC: minimum bactericidal concentration
<sup>4</sup>MIC<sub>50</sub>: MICs required to inhibit the growth of 50% of organisms
<sup>5</sup>MIC<sub>90</sub>: MICs required to inhibit the growth of 90% of organisms
<sup>6</sup>MIC<sub>50</sub>: MBCs required to inhibit the growth of 50% of organisms
<sup>7</sup>MIC<sub>90</sub>: MBCs required to inhibit the growth of 90% of organisms

### Effect of pH, temperature, and ions on CHAP^SAP26^-161 activity

Figure 3 demonstrates the effects of temperature, pH, and various ions on the enzymatic activity of CHAP^SAP26^-161. The antimicrobial activity of CHAP^SAP26^-161 against *A. baumannii* and *S. aureus* remained >95% of its activity after 1 h of incubation at 4°C–37°C; however, it decreased by approximately 15%–20% at 60°C (Figure 3A). The effect of pH on the antibacterial activity of CHAP^SAP26^-161 against *A. baumannii* and *S. aureus* varied. The antimicrobial activity of CHAP^SAP26^-161 against *A. baumannii* remained unaffected by pH, but that against *S. aureus* was reduced by approximately 40–50% at pH 3 and 9–10 (Figure 3B). Interestingly, the addition of ZnCl_2_ dramatically increased the antibacterial activity of CHAP^SAP26^-161 compared with that with other divalent ions (Figure 3C).

**Figure 3.**
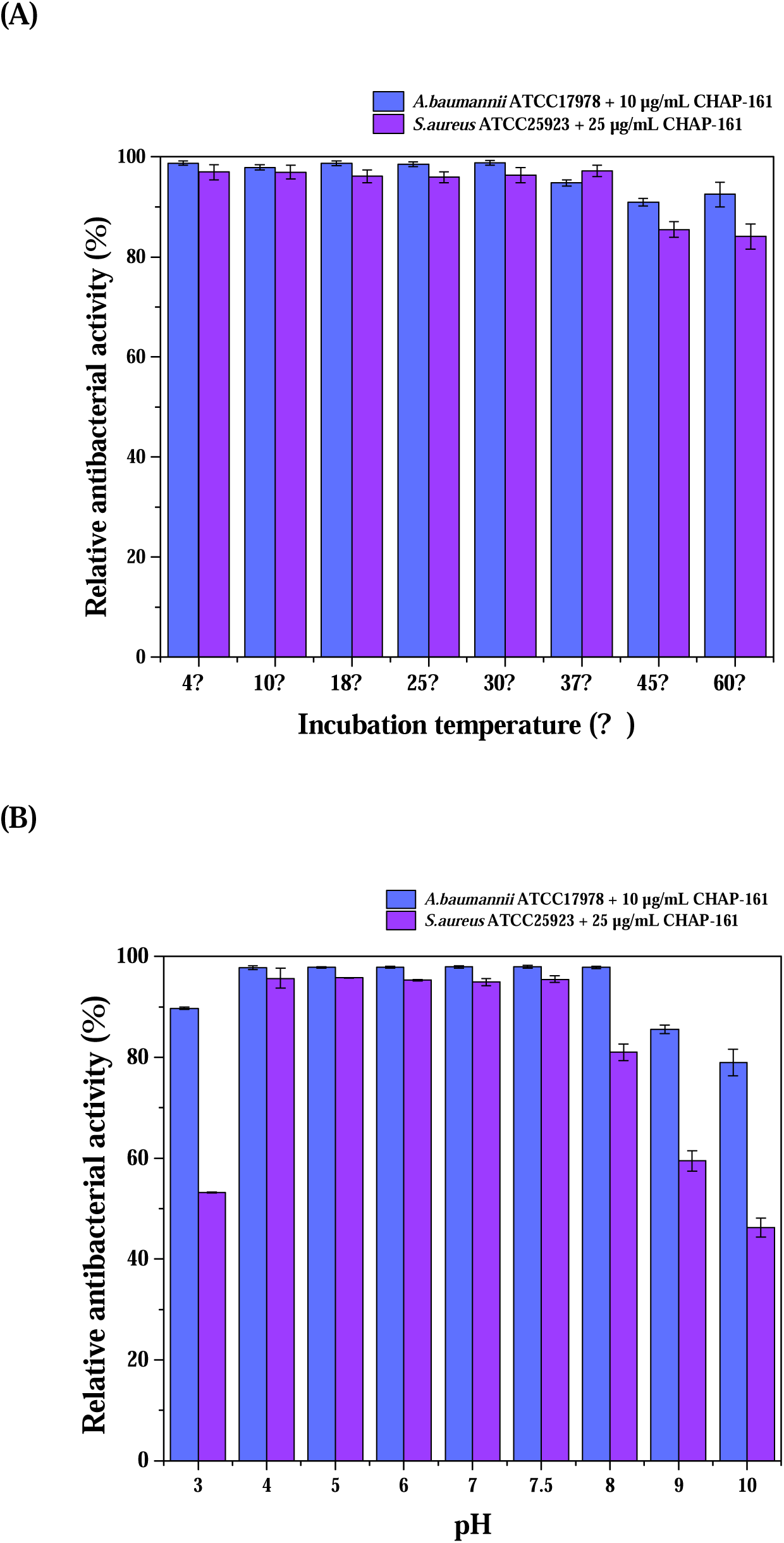
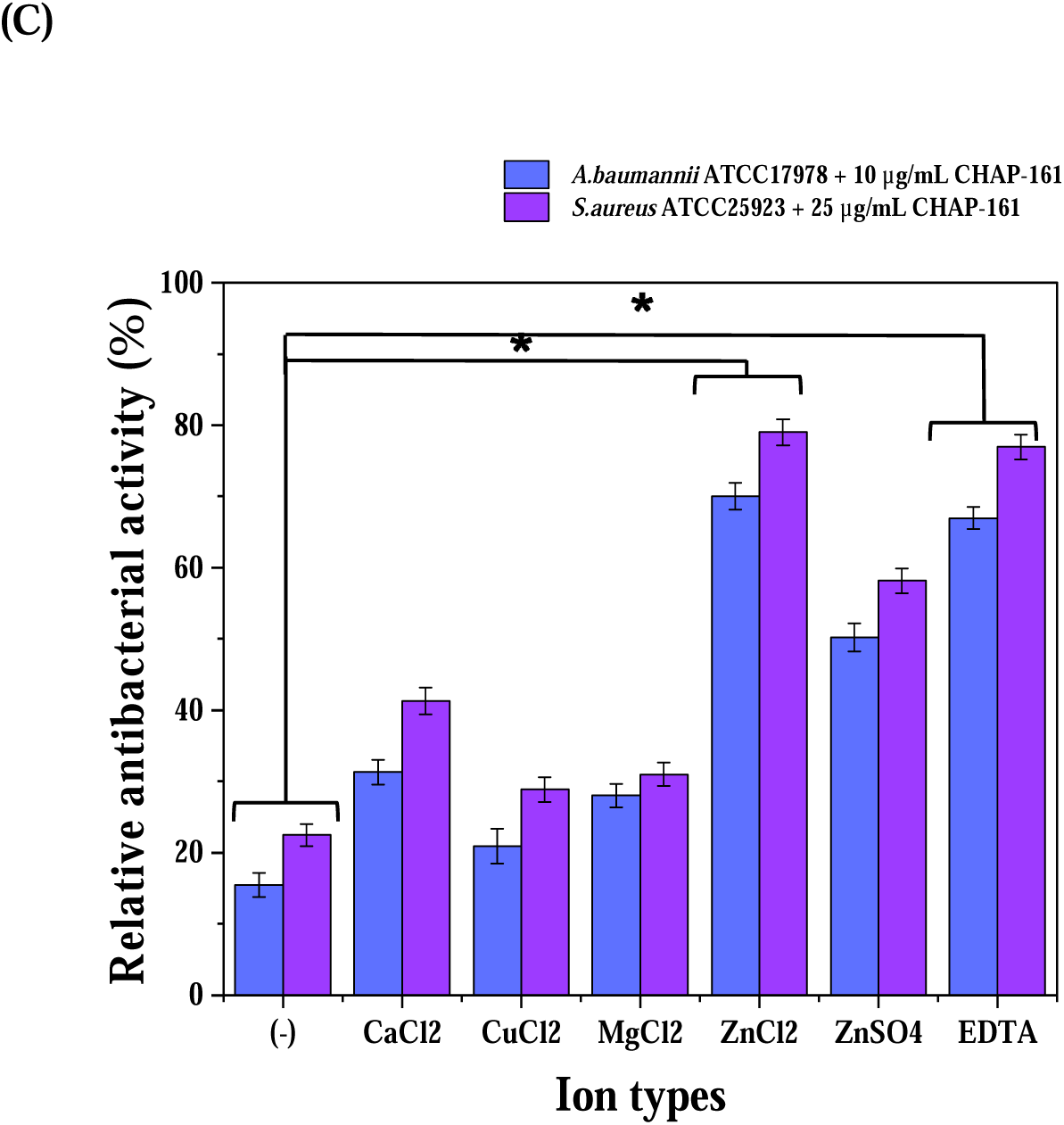
Impact of environmental conditions on the bactericidal activity of endolysins. Includes analysis of LysSAP26 (25 µg/mL for *A. baumannii*, 50 µg/mL for *S. aureus*) and CHAP^SAP26^-161 (5 µg/mL for *A. baumannii*, 10 µg/mL for *S. aureus*). (A) Bactericidal activity across a temperature range of 4°C to 60°C; (B) activity assessment across pH levels from 3 to 10; (C) assessing the bactericidal activity of endolysins in various ionic conditions, including NaCl (up to 500 mM) and divalent metal ions (1-mM each of CaCl_2_, CuCl_2_, MgCl_2_, ZnCl_2_, and ZnSO_4_), including EDTA concentrations up to 10 mM. \**p* < 0.001 showing statistical significance.

### Cytotoxicity of CHAP^SAP26^-161 and its efficacy in *A. baumannii* and *C. difficile-*infected mouse models

The MTT assay results showed that CHAP^SAP26^-161 did not affect the metabolic activity of A549 cells at concentrations of 25 μg/mL and 300 μg/mL, as evidenced by the over 90% cell survival rate. This indicates that CHAP^SAP26^-161 shows no cytotoxic effects on A549 cells up to a concentration of 500 μg/mL (refer to Supplementary Figure 1).

Experiments on mice were conducted to test the antibacterial effectiveness of CHAP^SAP26^-161 against two bacterial strains: *A. baumannii* (Figure 4). In our systemic infection model, neutropenic mice infected with *A. baumannii* were treated with CHAP^SAP26^-161 and LysSAP26 or given PBS as a control (Group 3), as part of a safety test, and showed no fatalities, similar to the inactive control (Group 1) (Figure 4B). Mice treated with 50 μg of CHAP^SAP26^-161 had a 100% survival rate over 7 days. Conversely, all patients treated with the same dose of LysSAP26 (Group 4) died within 2 days. Although less effective than Group 5, this result was improved over the infection control group (Group 2), where mice died just 1 day post infection.

**Figure 4.**
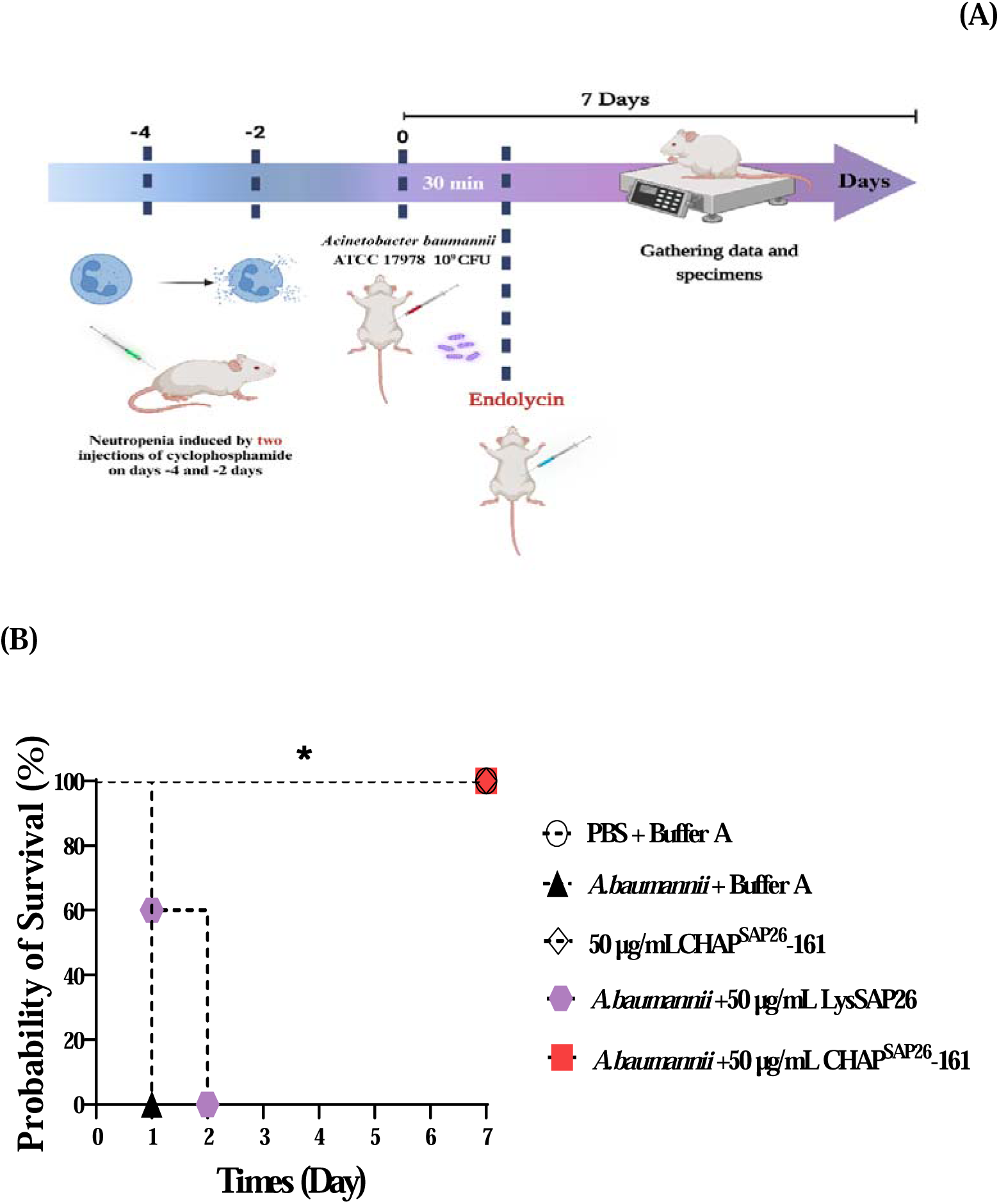
*In vivo* efficacy of CHAP-161 endolysin in mouse models infected with *Acinetobacter baumannii* and *Clostridioides difficile*. Statistical significance was observed (*p < 0.01). (A) CHAP-161 (50 μg/200 μL) and LysSAP26 (50 μg/200 μL) tested in BALB/c mice infected with *A. baumannii*, with a 7-day monitoring period; (B) CHAP^SAP26^-161 (150 μg/200 μL and 300 μg/200 μL) and LysSAP26 (300 μg/200 μL) tested in C57BL/6 mice infected with *C. difficile*, monitored over 14 days.

## Discussion

CHAP domains, which are integral components of bacterial amidases, autolysins, and bacteriophage-encoded peptidoglycan hydrolases, have been identified as critical agents in cleaving the molecular bridges that link peptidoglycan strands within bacterial cell walls, thereby facilitating cell lysis (Vermassen et al., 2019). These domains are characteristic of a broader family of murein hydrolases, distinguished by Cys and His residues within their active sites, which are critical for their catalytic activity (Abram et al., 2020). LysK, a truncated endolysin featuring a CHAP domain, consists of 495 amino acid residues. This endolysin is characterized by a CHAP domain spanning residues 35 to 160, an amidase-2 domain extending from residues 197 to 346, and an SH3b domain between residues 412 and 481 Horgan et al., 2009a).

Horgan et al. showed that a truncated mutant of LysK (Horgan et al., 2009a), with deletions from residue 163 to the protein’s C-terminus, retained its lytic activity against methicillin-resistant *S. aureus* (MDR *S. aureus*). Two truncated mutants of LysSAP26, CHAP^SAP26^-139 and CHAP^SAP26^-161, were engineered by removing the C-terminal domain to evaluate their role as a potential cell binding module. The purification processes for CHAP^SAP26^-139 and CHAP^SAP26^-161 demonstrated superior outcomes compared with LysSAP26, with significant reductions in undesirable protein contamination and enhancements in protein yields at the final purification stage. Notably, CHAP^SAP26^-161 exhibited the highest yield among the proteins tested, suggesting that the absence of the C-terminal domain contributes to its solubility in the purification buffer. Moreover, the purity of CHAP^SAP26^-161 was further enhanced via repeated affinity or size exclusion chromatography. Among the two truncated LysSAP26 mutants, CHAP^SAP26^-139 showed limited or weaker activity against all tested bacterial species than the wild-type and CHAP^SAP26^-161 proteins. These findings indicate that the amino acid residues from 140 to 161 are crucial for the enzymatic activity of CHAP. Notably, CHAP^SAP26^-161 demonstrated bactericidal activities that were 2.5–5-fold greater against *S. aureus* and *A. baumannii* than the wild-type protein, according to protein weight. Altogether, we produced a smaller CHAP-containing protein possessing similar or better antimicrobial activity and protein purification yields than LysSAP26.

The phage Twort endolysin (PlyTW) is composed of three distinct domains: a CHAP domain, an amidase-2 domain, and an SH3b-5 cell binding domain (CBD) (Becker et al., 2015). Notably, the isolated CHAP domain is capable of lysing *S. aureus in vivo*, with the absence of CBD resulting in a 10-fold decline in enzymatic efficacy. Yu et al. engineered a truncated variant of LysSAP33 (residues 1–156; CHAP-156), which shares identity with LysSAP26, albeit originating from a different phage, and observed a significant diminution in lytic efficiency upon removal of the C-terminal domain, underscoring its importance (Yu et al., 2021). Contrary to the findings of Yu et al., our research posits that CHAP proteins lacking the C-terminal domain, exemplified by CHAP^SAP26^-161, show superior antimicrobial prowess against a broader spectrum of bacterial strains compared with the native enzyme. In addition, our results in conjunction with those of Yu et al. suggest that the segment comprising residues 1–161 of LysSAP26 represents the minimal functional unit capable of achieving optimal antimicrobial activity upon C-terminal truncation.

CHAP^SAP26^-161 exhibited antimicrobial activities that were nearly comparable to those of LysSAP26, albeit under conditions of strong acidity and alkalinity. This indicates that the C-terminal domain may be crucial in maintaining protein stability under such extreme pH conditions. To enhance enzymatic function and demonstrate an efficient bactericidal effect upon the addition of EDTA, both the wild-type and CHAP^SAP26^-161 proteins required the presence of zinc cations. Conversely, PlyTW demonstrated increased antibacterial activity in the presence of Ca^2+^ ions but inhibited activity when exposed to zinc ions or EDTA (Becker et al., 2015).

Our in vivo experiments conducted on mice to evaluate the antibacterial efficacy of CHAP^SAP26^-161 against *A. baumannii* suggest that CHAP^SAP26^-161 possesses considerable therapeutic potential (Figure 4). In a neutropenic mouse model infected with *A. baumannii*, treatment with CHAP^SAP26^-161 resulted in a 100% survival rate, significantly improving compared with the LysSAP26 treatment group (Figure 4 B). Notably, the survival time in the CHAP^SAP26^-161-treated group was substantially enhanced relative to that in the infection control group. This study provides a deeper understanding of the therapeutic effects of CHAP^SAP26^-161 and offers crucial foundational data for future research on developing antimicrobial peptides.

Considering the use of these proteins for clinical or veterinary treatments, the aforementioned ionic conditions must be considered. Preliminary findings from our pilot study indicate that CHAP^SAP26^-161 has potential as an effective antimicrobial agent against pathogens such as *Bacillus megaterium*, *Bacillus muralis*, *Corynebacterium striatum*, and *E. faecium,* which are known to contaminate catheters in clinical environments (Choi et al., 2022).

In summary, our investigation has led to the development of a truncated CHAP-containing protein variant, CHAP^SAP26^-161, which exhibits antimicrobial activity and protein purification yields at par with or exceeding those of wild-type LysSAP26. To the best of our knowledge, this is the first report on the CHAP^SAP26^-161 protein demonstrating antibacterial activity against *C. difficile*, in addition to its efficacy against methicillin-resistant *S. aureus* (MDR *S. aureus*) and carbapenem-resistant *A. baumannii*. Given the significant challenge in generating resistant mutants to endolysins, CHAP^SAP26^-161 has emerged as a potential antibacterial candidate for combating drug-resistant bacterial infections.

## Figure Legends

**Supplementary Figure 1.**
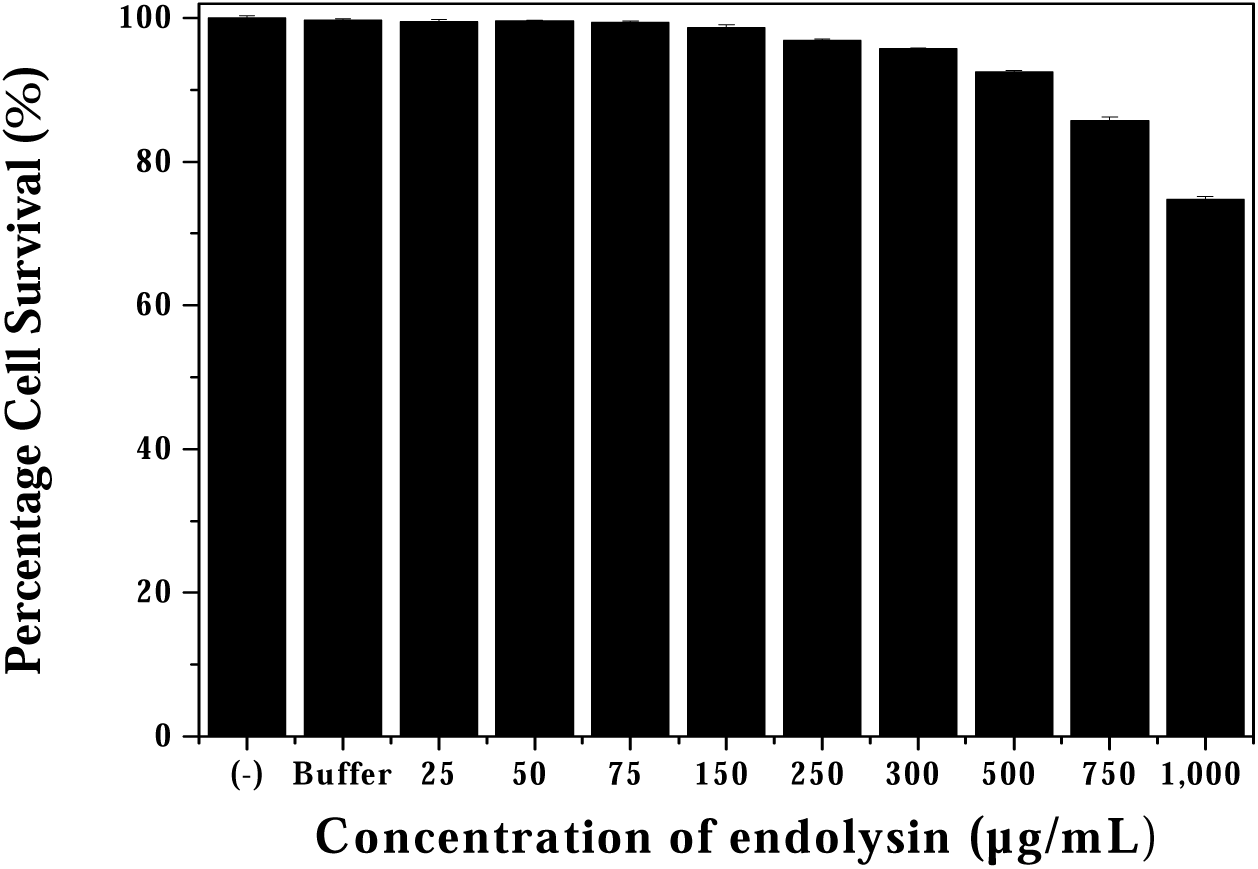
Cytotoxicity assessment of CHAPSAP26-161 on human pulmonary epithelial cells. The cytotoxicity assessment of CHAPSAP26-161 on A549 cells using the MTT assay within a concentration range of 25 to 1000 μg/mL is demonstrated.

**Supplementary Table 1.**
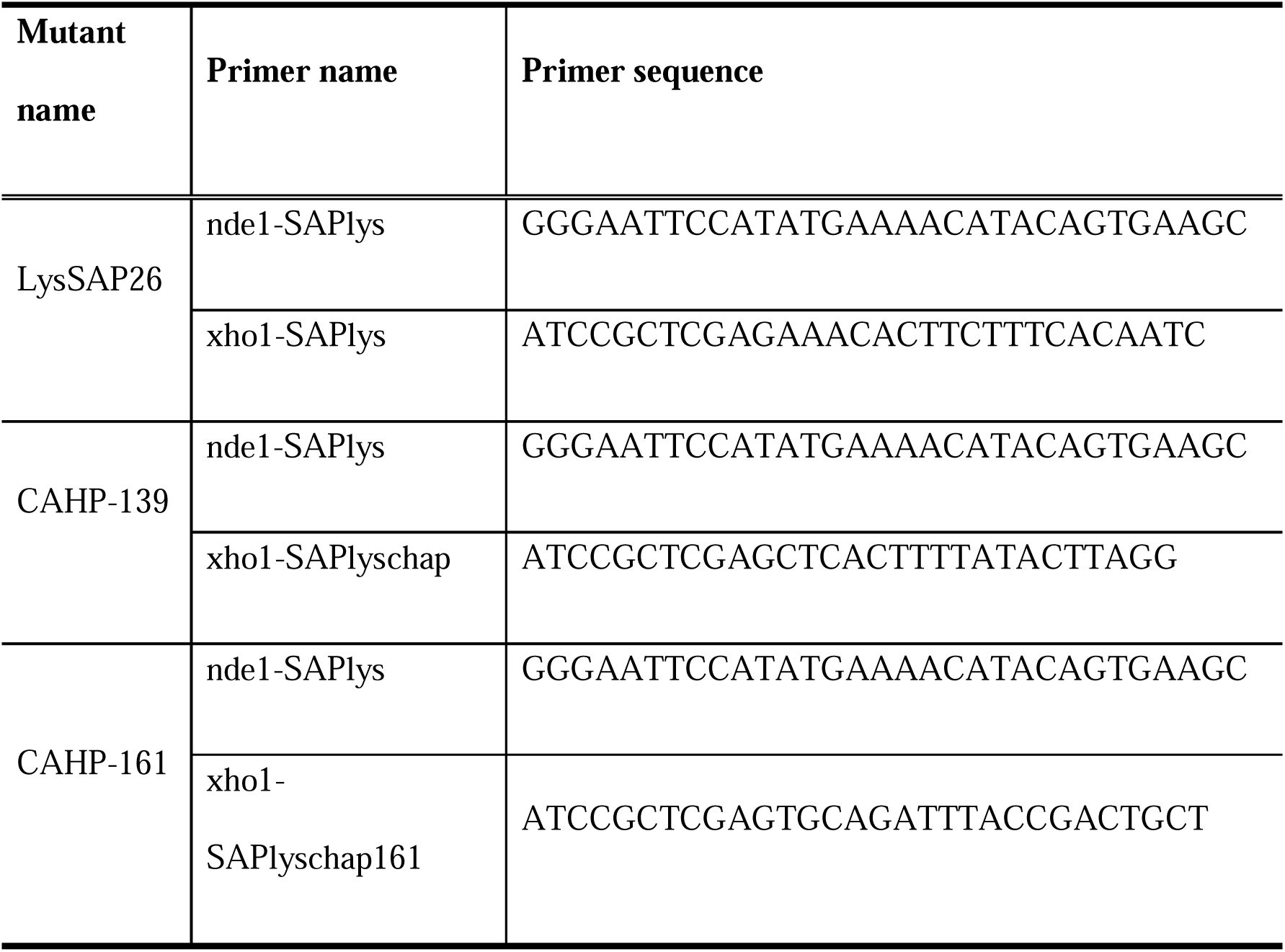
Type and sequence of primers used.

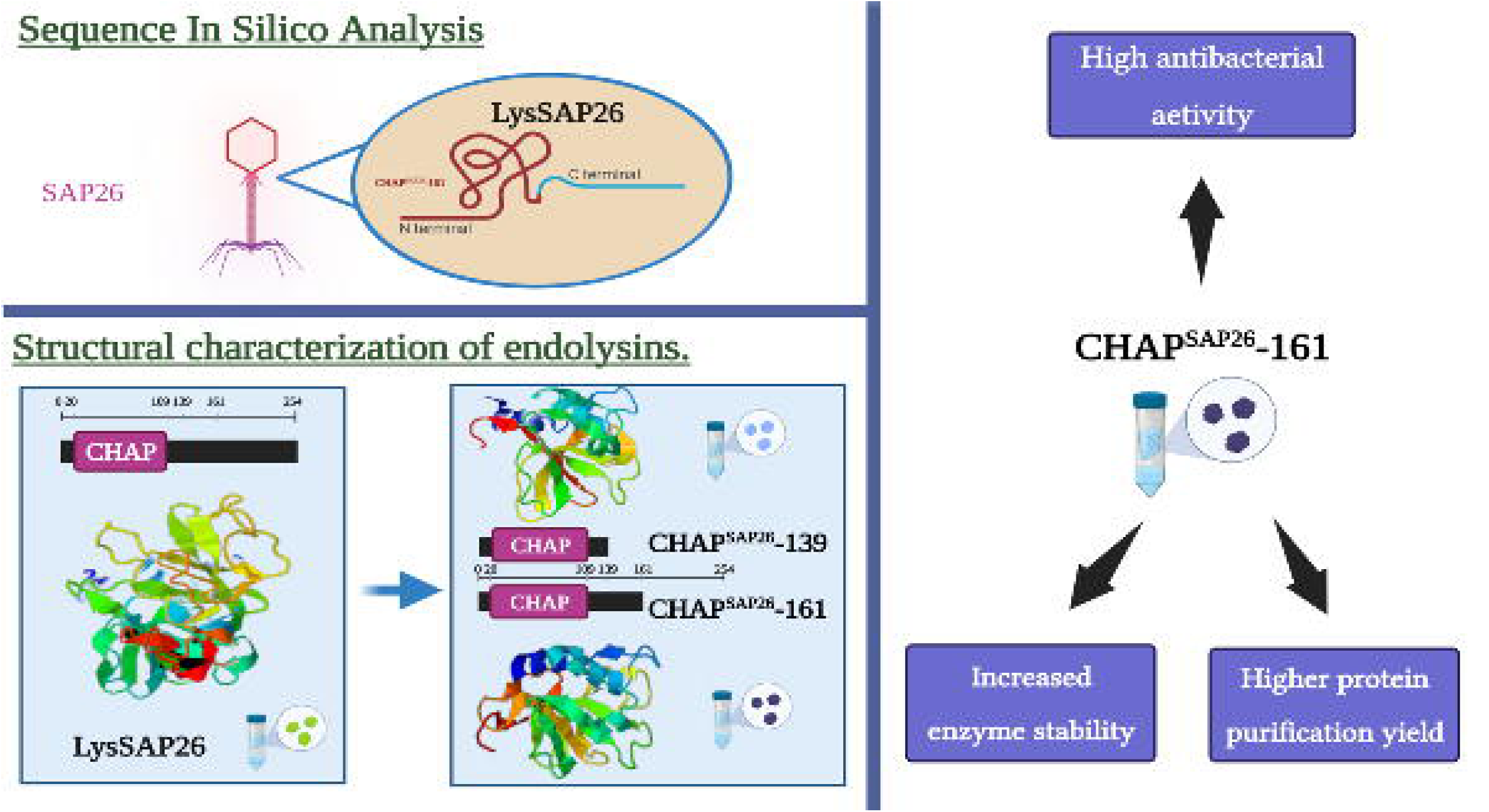

## References

1. Abdelrahman, F., Easwaran, M., Daramola, O.I., Ragab, S., Lynch, S., Oduselu, T.J., Khan, F.M., Ayobami, A., Adnan, F., Torrents, E., Sanmukh, S., El-Shibiny, A., 2021. Phage-encoded endolysins. Antibiotics (Basel). 10, 124.

2. Abram, T.J., Cherukury, H., Ou, C.Y., Vu, T., Toledano, M., Li, Y., Grunwald, J.T., Toosky, M.N., Tifrea, D.F., Slepenkin, A., Chong, J., Kong, L., Del Pozo, D.V., La, K.T., Labanieh, L., Zimak, J., Shen, B., Huang, S.S., Gratton, E., Peterson, E.M., Zhao, W., 2020. Rapid bacterial detection and antibiotic susceptibility testing in whole blood using one-step, high throughput blood digital PCR. Lab Chip. 20, 477–489.

3. Bateman, A., Rawlings, N.D., 2003. The CHAP domain: a large family of amidases including GSP amidase and peptidoglycan hydrolases. Trends Biochem Sci. 28, 234–7.

4. Becker, S.C., Swift, S., Korobova, O., Schischkova, N., Kopylov, P., Donovan, D.M., Abaev, I., 2015. Lytic activity of the staphylolytic Twort phage endolysin CHAP domain is enhanced by the SH3b cell wall binding domain. FEMS Microbiol Lett. 362, 1–8.

5. Boneca, I.G., 2021. The future of microbial drug resistance. Microb Drug Resist. 27, 1–2.

6. Borysowski, J., Weber-DaLbrowska, B., Górski, A., 2006. Bacteriophage endolysins as a novel class of antibacterial agents. Exp Biol Med (Maywood). 231, 366–77.

7. Choi, Y.J., Kim, S., Bae, S., Kim, Y., Chang, H.H., Kim, J., 2022. Antibacterial effects of recombinant endolysins in disinfecting medical equipment: a pilot study. Front Microbiol. 12:773640.

8. CLSI, 2020. Performance standards for antimicrobial susceptibility testing 1–352.

9. CLSI GUIDELINES 2020. URL https://www.pdffiller.com/jsfiller-desk10/?projectId=61a5afdc300a5a02ea1471d0&lp=true#38e05859d4144a42bd0e11c500c80f19.

10. Nelson DC, Schmelcher M, Rodriguez-Rubio L, Klumpp J, Pritchard DG, Dong S, Donovan DM., 2012. Endolysins as antimicrobials. Adv Virus Res. 83, 299–365.

11. Fenton, M., Ross, R.P., Mcauliffe, O., O’Mahony, J., Coffey, A., 2011. Characterization of the *Staphylococcal* bacteriophage lysin CHAP K. J Appl Microbiol. 111, 1025–1035.

12. Filatova, L.Y., Becker, S.C., Donovan, D.M., Gladilin, A.K., Klyachko, N.L., 2010. LysK, the enzyme lysing *Staphylococcus aureus* cells: Specific kinetic features and approaches towards stabilization. Biochimie. 92, 507–513.

13. Fujimoto, K., Uematsu, S., 2022. Phage therapy for *Clostridioides difficile* infection. Front Immunol. 13, 1057892.

14. Gondil, V.S., Harjai, K., Chhibber, S., 2020. Endolysins as emerging alternative therapeutic agents to counter drug-resistant infections. Int J Antimicrob Agents. 55.

15. Hecke, T. Van, 2013. Power study of ANOVA versus Kruskal-Wallis test. 10.1080/09720510.2012.10701623. 15, 241–247.

16. Heselpoth, R.D., Swift, S.M., Linden, S.B., Mitchell, M.S., Nelson, D.C., 2021. Enzybiotics: endolysins and bacteriocins. Bacteriophages. 989–1030. 34.

17. Horgan, M., O’Flynn, G., Garry, J., Cooney, J., Coffey, A., Fitzgerald, G.F., Paul Ross, R., McAuliffe, O., 2009b. Phage lysin LysK can be truncated to its CHAP domain and retain lytic activity against live antibiotic-resistant *Staphylococci*. Appl Environ Microbiol. 75, 872–874.

18. Khan, A., Joshi, H., 2022. Simple Two-step, High yield protocol for isolation and amplification of bacteriophages against methicillin-resistant Staphylococcus aureus (MRSA). Curr Protoc. 2, e395.

19. Kim, K., Islam, M.M., Kim, D., Yun, S.H., Kim, J., Lee, J.C., Shin, M., 2021. Characterization of a novel phage ΦAb1656-2 and its endolysin with higher antimicrobial activity against multidrug-resistant *Acinetobacter baumannii*. Viruses. 13, 1848.

20. Kim, S., Jin, J., Lee, D., Kim, J., 2020a. Antibacterial activities of and biofilm removal by Ablysin, an endogenous lysozyme-like protein originated from *Acinetobacter baumannii* 1656-2. J Glob Antimicrob Resist. 23, 297–302.

21. Kim, S., Jin, J.-S., Choi, Y.-J., Kim, J., 2020b. LysSAP26, a new recombinant phage endolysin with a broad-spectrum antibacterial activity. Viruses. 12, 1340.

22. Kim, S., Lee, D., Jin, J., Kim, J., 2020c. Antimicrobial activity of LysSS, a novel phage endolysin, against *Acinetobacter baumannii* and *Pseudomonas aeruginosa*. J Glob Antimicrob Resist. 22, 32–39.

23. Kortright, K.E., Chan, B.K., Koff, J.L., Turner, P.E., 2019. Phage Therapy: a renewed approach to combat antibiotic-resistant bacteria. Cell Host Microbe. 25, 219–232.

24. Kretzer, J.W., Lehmann, R., Schmelcher, M., Banz, M., Kim, K.P., Korn, C., Loessner, M.J., 2007. Use of high-affinity cell wall-binding domains of bacteriophage endolysins for immobilization and separation of bacterial cells. Appl Environ Microbiol. 73, 1992–2000.

25. Mc, J., Av, F., Fk, Z., Frp, B., Mr, M., Rg, M., Pds, N., 2021. Multidrug-resistant hospital bacteria: epidemiological factors and susceptibility profile. Microb Drug Resist. 27, 433–440.

26. McHugh, M.L., 2011. Multiple comparison analysis testing in ANOVA. Biochem Med (Zagreb). 21, 203–209.

27. Mirski, T., Mizak, L., Nakonieczna, A., Gryko, R., 2019. Bacteriophages, phage endolysins and antimicrobial peptides – The possibilities for their common use to combat infections and in the design of new drugs. Ann Agric Environ Med. 26, 203–209.

28. Mondal, S.I., Akter, A., Draper, L.A., Ross, R.P., Hill, C., 2021. Characterization of an endolysin targeting *Clostridioides difficile* that affects spore outgrowth. Int J Mol Sci. 22, 8690

29. Mondal, S.I., Draper, L.A., Ross, R.P., Hill, C., 2020. Bacteriophage endolysins as a potential weapon to combat *Clostridioides difficile* infection. Gut Microbes. 12, 1813533.

30. Murdoch, D.J., Tsai, Y.L., Adcock, J., 2012. P-values are random variables. 62, 242–245.

31. Nelson, D.C., Schmelcher, M., Rodriguez-Rubio, L., Klumpp, J., Pritchard, D.G., Dong, S., Donovan, D.M., 2012. Adv Virus Res. 83, 299–365.

32. Noor, A., Khetarpal, S., 2023. Anaerobic Infections. StatPearls.

33. O’Flaherty, S., Coffey, A., Meaney, W., Fitzgerald, G.F., Ross, R.P., 2005. The recombinant phage lysin LysK has a broad spectrum oflytic activity against clinically relevant *Staphylococci*, including methicillin-resistant *Staphylococcus aureus*. J Bacteriol, 187, 7161.

34. Ozma, M.A., Abbasi, A., Asgharzadeh, M., Pagliano, P., Guarino, A., Köse, S., Kafil, H.S., 2022. Antibiotic therapy for pan-drug-resistant infections. Infez Med, 30, 525–531.

35. Sandhu, B.K., McBride, S.M., 2018. Clostridioides difficile. Trends Microbiol, 26, 1049–1050.

36. Schmelcher, M., Donovan, D.M., Loessner, M.J., 2012. Bacteriophage endolysins as novel antimicrobials. Future Microbiol, 7, 1147–1171.

37. Schmelcher, M., Loessner, M.J., 2016. Bacteriophage endolysins: applications for food safety. Curr Opin Biotechnol, 37, 76–87.

38. Sekiya, H., Yamaji, H., Yoshida, A., Matsunami, R., Kamitori, S., Tamai, E., 2022. Biochemical characterizations of the putative endolysin Ecd09610 catalytic domain from *Clostridioides difficile*. Antibiotics (Basel), 11, 1131.

39. Souza, S.G.P. De, Santos, I.C. Dos, Bondezan, M.A.D., Corsatto, L.F.M., Caetano, I.C.D.S., Zaniolo, M.M., Matta, R. Da, Merlini, L.S., Barbosa, L.N., Gonçalves, D.Di., 2021. Bacteria with a potential for multidrug resistance in hospital material. Microb Drug Resist 27, 835–842.

40. Stacy, A., Andrade-Oliveira, V., McCulloch, J.A., Hild, B., Oh, J.H., Perez-Chaparro, P.J., Sim, C.K., Lim, A.I., Link, V.M., Enamorado, M., Trinchieri, G., Segre, J.A., Rehermann, B., Belkaid, Y., 2021. Infection trains the host for microbiota-enhanced resistance to pathogens. Cell. 184, 615–627.e17.

41. Stone, E., Campbell, K., Grant, I., McAuliffe, O., 2019. Understanding and exploiting phage–host interactions. Viruses. 11, 567–567.

42. Sundarrajan, S., Raghupatil, J., Vipra, A., Narasimhaswamy, N., Saravanan, S., Appaiah, C., Poonacha, N., Desai, S., Nair, S., Bhatt, R.N., Roy, P., Chikkamadaiah, R., Durgaiah, M., iram, B., Padmanabhan, S., Sharma, U., 2014. Bacteriophage-derived CHAP domain protein, P128, kills Staphylococcus cells by cleaving interpeptide cross-bridge of peptidoglycan. Microbiology (United Kingdom). 160, 2157–2169.

43. Ul Haq, I., Chaudhry, W.N., Akhtar, M.N., Andleeb, S., Qadri, I., 2012. Bacteriophages and their implications on future biotechnology: A review. Virol J. 9, 1–8.

44. Vermassen, A., Leroy, S., Talon, R., Provot, C., Popowska, M., Desvaux, M., 2019. Cell wall hydrolases in bacteria: Insight on the diversity of cell wall amidases, glycosidases and peptidases toward peptidoglycan. Front Microbiol. 10, 331.

45. Vollmer, W., Blanot, D., De Pedro, M.A., 2008. Peptidoglycan structure and architecture. FEMS Microbiol Rev. 32, 149–67.

46. Yu, J.H., Park, D.W., Lim, J.A., Park, J.H., 2021. Characterization of *Staphylococcal* endolysin LysSAP33 possessing untypical domain composition. Journal of Microbiology. 59, 840–847.

